# RNA-Seq analysis reveals pluripotency-associated genes and their interaction networks in human embryonic stem cells

**DOI:** 10.1101/782169

**Authors:** Arindam Ghosh, Anup Som

**Affiliations:** Centre of Bioinformatics, Institute of Interdisciplinary Studies, University of Allahabad

**Keywords:** Embryonic stem cells, RNA-seq, Differential gene expression, Critical genes, Co-expression network, GO enrichment, Pathway analysis

## Abstract

Insight into the key genes of pluripotency in human and their interrelationships is necessary for understanding the underlying mechanism of pluripotency and hence their successful application in regenerative medicine. The recent advances in transcriptomics technologies have created new opportunities to decipher the genes involved in pluripotency, genetic network that governs the unique properties of embryonic stem cells and lineage differentiation mechanisms in a deeper scale. There are a large number of experimental studies on human embryonic stem cells (hESCs) being routinely conducted for unfolding the underlying biology of embryogenesis and their clinical prospects. However, the outcome of these studies often lacks consensus due to differences in samples, experimental techniques and/or analysis protocols. A universal stemness gene list is still lacking. In this quest, we compared transcriptomic profiles of pluripotent and non-pluripotent samples from diverse cell lines/types generated through RNA-sequencing (RNA-seq). We used a uniform pipeline for the analysis of raw RNA-seq data in order to reduce the amount of variation. Our analysis revealed a consensus set of 498 pluripotency-associated genes and 432 genes as potential pluripotent cell differentiation markers. Furthermore, we predicted 32 genes as “pluripotency critical genes”. Reconstruction and analysis of co-expression networks further highlighted the importance of these genes. Gene ontology (GO) and pathway enrichment analysis, StemChecker and literature survey confirmed the involvement of the genes in the induction and maintenance of pluripotency, though more experimental studies are required for understanding their molecular mechanisms in human.

## 1. INTRODUCTION

Pluripotency is the ability of a cell to develop into the three primary germ cell layers (i.e., endoderm, mesoderm and ectoderm) of the early embryo and therefore into all cells of the adult body, but not extra-embryonic tissues such as the placenta. Embryonic stem cells (ESCs) are pluripotent cells. The ability to give rise to any mature cell type makes the pluripotent stem cells as an ideal cell candidate for regenerative medicine (De Los Angeles et al., 2015). However successful application of pluripotent stem cells is hampered due to lack of complete knowledge of the genes/proteins involved in pluripotency and their underlining mechanisms. The molecular mechanisms behind this process are complex interplay between the genes/proteins that includes transcription factors with epigenetic regulators and signaling proteins. Therefore, identification of genes/proteins involved in pluripotency and their interrelationships is necessary for understanding the induction/loss and maintenance of pluripotency.

Ever since the first isolation of human embryonic stem cells (hESCs), numerous studies have been conducted to identify the essential genes and their underlying coordinated expression mechanisms that are involved in the establishment and maintenance of pluripotency. Earlier Microarray technology provided a unique opportunity to examine gene expression patterns, thus a good number of studies on hESC were carried out based on differentially expressed genes between pluripotent and non-pluripotent samples using microarray gene expression data. For example, Bhattacharya et al (2004) compared expression profiles of hESC lines against human universal RNA and identified a common subset of 92 genes that constitute molecular signature of hESCs. In other work, Muller et al (2008) used unsupervised machine learning approach, and identified 299 genes and uncovered a possible molecular network describing human pluripotency (called PluriNet). Another similar work was based on machine learning approach where Newman & Cooper (2010) developed a method for unsupervised clustering called AutoSOME and identified a cluster of 3421 genes upregulated in pluripotency (called PluriUP). They further mapped these genes to the Human Protein Reference Database (HPRD) and extracted a network (called PluriPlus) consisting of a subset of 1165 PluriUp genes and 2463 protein-protein interactions (PPIs). Chia et al (2010) carried out whole genome RNA interference (RNAi) screening targeting 21,121 human genes and identified 566 candidate genes which regulate self-renewal and pluripotency properties in hESCs. Som et al (2012) derived a putative human pluripotency network from a mouse model network (Som et al., 2010) that consist of 136 genes/proteins and 196 interactions. Assou et al (2007) performed a meta-analysis of 38 original studies reporting transcriptome of hESCs. Their study revealed only one consensus hESC gene (POU5F1) supported by all studies, 1076 overexpressed genes supported by at least three independent studies and similarly 783 downregulated genes reported in at least three studies. In a recent work, Yilmaz et al (2018) used CRISPR-Cas9 based genome-wide screens and identified a set of 50 genes essential for the normal growth and survival of human pluripotent stem cells. Besides the computational approaches, Narad et al (2017) assembled a literature curated network of 122 genes/proteins 166 molecular interactions depicting the cellular and molecular mechanisms encompassing pluripotency (called hPluriNet). However comparison with previous studies revealed a lack of consensus of the essential genes required in the induction and maintenance of pluripotency in human.

Over the last decade, RNA-sequencing (RNA-seq) has evolved as a preferred method for transcriptomic profiling. Unlike microarray technology that suffers from two major limitations of the lack of common set of probes in the microarray chips and the availability of raw data for reanalysis, it does not require prior information of the organism’s genome for measuring gene expression and additionally it is able to detect novel transcripts and isoforms, and genes with very low expression levels (Zhao et al., 2014). RNA-seq has been extensively used to profile hESC transcriptome and also to study the changes occurring during its differentiation (Wu et al., 2010; Sun et al., 2018). In this study, we used computational approach for identifying the pluripotency-associated genes based on consensus method. Our analysis of RNA-seq based transcriptome profiles of pluripotent and non-pluripotent samples from five different datasets revealed 498 “pluripotency-associated genes” including 32 genes critical for pluripotency and 432 genes as potential pluripotent cell differential markers. Reconstruction and analysis of co-expression network depicted the pluripotency critical genes to form a small world network, thus allowing for efficient transfer of information for maintenance of pluripotency. Validations of the identified genes were carried out through gene ontology (GO) and pathway enrichment analysis, StemChecker (Pinto et al., 2015) and literature survey.

## 2. MATERIALS AND METHODS

### 2.1. Data retrieval

Raw RNA-seq data in fastq format for five datasets were retrieved from the European Nucleotide Archive (ENA) (Leinonen et al., 2010). We considered the data where each dataset contained transcriptome profiles of both pluripotent hESC and non-pluripotent samples (either derived from hESC or isolated from somatic cell). Table 1 provides a summary of the raw data included in our study, while Supplementary File 1 lists all the sample accession numbers.

**Table 1:** Summary of raw data included in the study.

| Dataset | Project ID | Number of pluripotent samples | Number of non-pluripotent samples | Sequencer | Sequencing type | References |
| --- | --- | --- | --- | --- | --- | --- |
| 1 | PRJNA307622 | 2 | 2 | Illumina HiSeq 2500 | Paired | Tompkins et al., 2016 |
| 2 | PRJNA338181 | 4 | 4 | Illumina HiSeq 2000 | Paired | Liu et al., 2017 |
| 3 | PRJNA296379 | 35 | 10 | Illumina HiSeq 2000 | Paired | Choi et al., 2015 |
| 4 | PRJNA230824 | 4 | 2 | Illumina HiSeq 2000 | Paired | Ma et al., 2014 |
| 5 | PRJNA422061 | 3 | 3 | Illumina NextSeq 500 | Single | Yilmaz et al., 2014 |

### 2.2. Data analysis

The quality of the raw reads were assessed using FastQC v0.11.5 toolkit (Andrews, 2010). Adapters and low quality reads were trimmed using Trimmomatic v0.36 (Bolger et al., 2014). HiSat2 v2.1.0 (Kim et al., 2015) was then used for mapping the raw reads to the reference genome GRCh38.p5 (Ensembl release 84). The expression for each gene were evaluated using featureCounts v1.6.2 (Liao et al., 2013) based on annotation from Ensembl release 84. The low expressed genes which did not have more than one count per million reads (1CPM) in at least two samples within each datasets were removed from further analysis. The raw counts were then normalized and used for differential expression testing using DESeq2 (Love et al., 2014). Additionally, edgeR (Robinson et al., 2010) and limma (Ritchie et al., 2015) were also used to validate the results of DESeq2. The genes with log_2_fc > 1.0 and adjusted p-value (p_adj_) < 0.01 were considered upregulated and overexpressed in pluripotency while those with log_2_fc < -1.0 and p_adj_ < 0.01 were considered downregulated or underexpressed in pluripotency. The entire workflow used in the study is shown in Figure 1.

**Figure 1:**
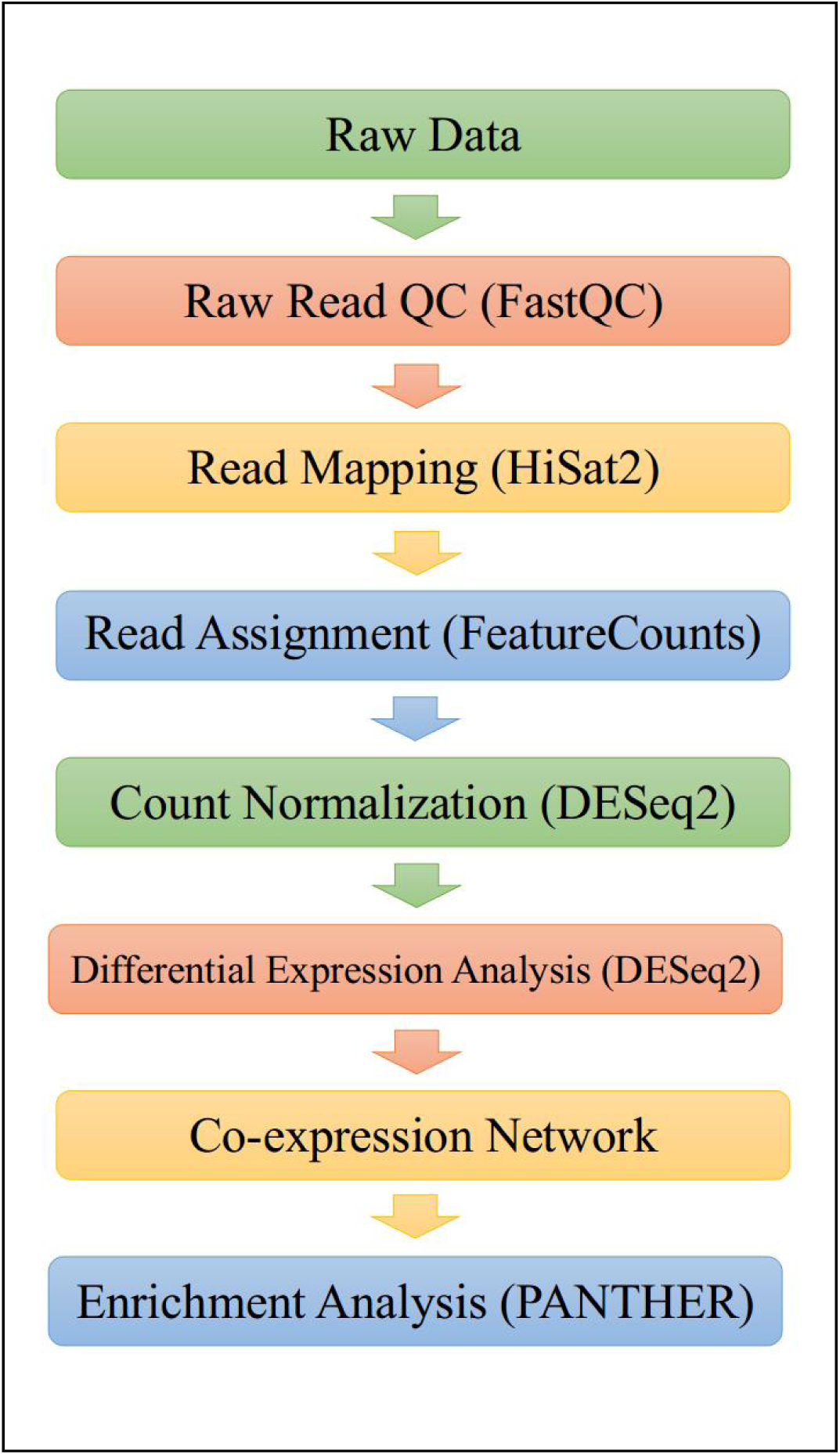
Overview of the workflow implemented in this study.

### 2.3. Co-expression network

Co-expression network was reconstructed separately for each dataset from the log transformed normalized count obtained from DESeq2. A threshold Pearson correlation value of 0.9 and Benjamini–Hochberg corrected p-value < 0.05 was set for two genes to be co-expressed. The networks were visualized using Cytoscape v3.6.1 (Shannon et al., 2003). Network Analyzer plugin of Cytoscape was used for calculating the topological parameters of the networks (Assenov et al., 2007). The networks were merged using the intersection mode in Cytoscape. Clustering of the merged network was done using the community cluster (GLay) algorithm in ClusterMaker (Morris et al., 2011). It is based on Newman-Girvan’s method of community clustering (Newman and Girvan, 2004; Su et al., 2010).

### 2.4. Enrichment Analysis

Identification of the enriched biological processes and pathways was carried out using PANTHER (Protein ANalysis THrough Evolutionary Relationships) (Mi et al., 2016). For biological processes, the GO Ontology database (Released 2019-02-02) and for pathways, Reactome version 65 (Released 2019-03-12) were used. Significance testing of the matched terms were performed using Fisher’s exact test. Significantly enriched terms were selected based on false discovery rate (FDR) adjusted p-value < 0.05.

## 3. RESULTS

### 3.1. Identification of genes specific to pluripotency

The objective of our study is to identify the genes and their underlying networks responsible for the induction and maintenance of human pluripotency. In order to achieve this, we compared the transcriptomic profiles of pluripotent and non-pluripotent samples using DESeq2. We also verified our list of overexpressed genes obtained from DESeq2 with two other popular methods of differential gene expression analysis namely edgeR and limma. The number of genes found to be dysregulated (upregulated/downregulated) by all the three tools indicated consensus in the detection of overexpressed genes for each dataset (Table 2). For further downstream analysis, only the consensus gene list from each dataset were considered. In all, we found 498 genes to be common across all the five datasets (Figure 2). This gene list will henceforth be referred to as the “pluripotency-associated gene (PAG)” list.

**Table 2:** Summary of dysregulated gene counts. The differentially expressed genes in each datasets were identified using three tools namely DESeq2, EdgeR and Limma. Here we list the number of upregulated and downregulated genes detected by each tool, the number of consensus genes detected by all the three tools in each dataset and the number of consensus genes across all the datasets.

| Dataset | Upregulated genes |  |  |  |  | Downregulated genes |  |  |  |  |
| --- | --- | --- | --- | --- | --- | --- | --- | --- | --- | --- |
|  | DESeq2 | EdgeR | Limma | No. of consensus genes in each dataset | Top 10% genes | DESeq2 | EdgeR | Limma | No. of consensus genes in each dataset | Top 10% genes |
| 1 | 3400 | 3632 | 3632 | 3399 | 340 | 3451 | 3446 | 3446 | 3412 | 341 |
| 2 | 3168 | 3408 | 3347 | 3120 | 312 | 4307 | 4198 | 3884 | 3875 | 388 |
| 3 | 4269 | 4069 | 4130 | 3909 | 391 | 2858 | 3054 | 2902 | 2756 | 276 |
| 4 | 2776 | 2785 | 2337 | 2293 | 229 | 2565 | 2642 | 1876 | 1867 | 187 |
| 5 | 3921 | 3911 | 3904 | 3887 | 389 | 3068 | 3127 | 3125 | 3067 | 307 |
|  | No. of consensus genes across all dataset |  |  | **498** | **32** | No. of consensus genes across all dataset |  |  | **432** | **8** |

**Figure 2:**
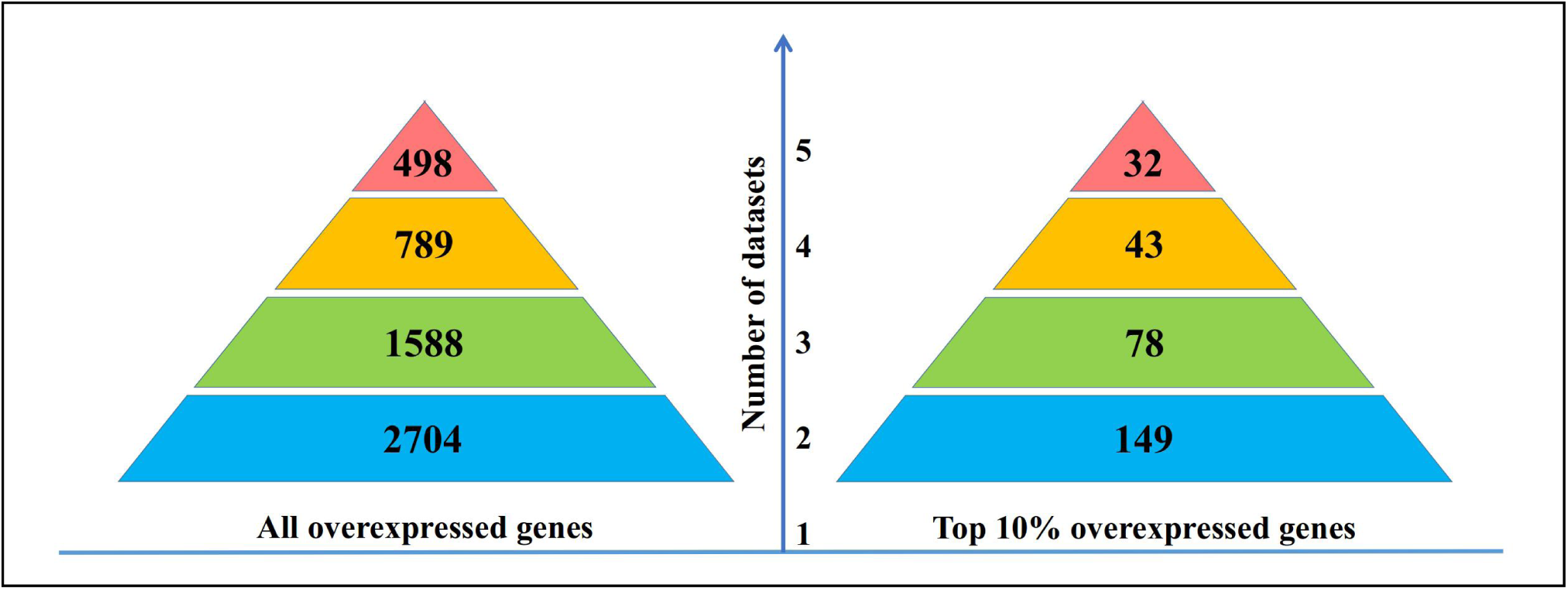
Identification of candidate genes based on the consensus of data. Analysis of the overexpressed genes in the five datasets revealed that as the number of dataset increases, the number of consensus genes decreases. A similar trend was observed when only the top 10% overexpressed genes in each dataset was considered.

We hypothesized that the genes those are critical for the induction and maintenance of pluripotency must be expressed at levels much higher than in non-pluripotent samples i.e., genes critical to pluripotency must have high fold-change. Thus, we took to study the top 10% positively dysregulated genes in each dataset. The fold-change values computed by DESeq2 were used for sorting the genes. We found 32 genes common among the most upregulated genes across all the datasets (Figure 2) and define them as “pluripotency critical genes (PCGs)”.

### 3.2. Co-expression network of pluripotency-associated genes

It is known that if genes are functionally related or involved in the same pathway or controlled by the same transcriptional regulatory program they are turned on or off at the same time (Weirauch, 2011). In a co-expression network, genes are linked if their expression levels are correlated. We built individual correlation networks of the pluripotency specific genes for each dataset and then merged them to keep only the overlapping nodes and edges among them. This merged network is a representation of the genes that were found to be correlated in all the five datasets and contained 490 PAGs. We refer to this as the PAG network and is provided as Supplementary File 2. Eight PAGs did not appear in this network and the possible explanation might be the fact that individual genes that are found to be differentially expressed need not be correlated. Methods for identification of differentially expressed genes test only individual genes rather than the effect on networked genes (Ahn et al., 2016; Irigoien and Arenas, 2018). Furthermore, the PAG network consisted of one major component of 476 nodes and 14 other island nodes. The occurrence of island nodes here further strengthens the fact that though these nodes/genes were overexpressed in all the datasets, their correlation could not be detected based on the considered threshold.

Network of the “pluripotency critical genes” (or PCG network) was extracted from the PAG network (Figure 3). The nodes of this network were found to be highly connected to each other having a density of 0.905, much higher than the PAG network (node density = 0.276). Other network topology parameters are listed in Table 3. The high clustering coefficient coupled with smaller characteristic path length indicates the PCG network to have a small-world architecture. This is not surprising for the nodes that were among the most overexpressed genes in each dataset. Nodes forming small world network are associated with important biological consequences as this level of connectivity allows for efficient and quick flow of signals within the network (Liao et al., 2017). The small world property thus highlights the importance of the 32 PCGs.

**Table 3:** Topological parameters of the co-expression networks of over-expressed genes.

| **Network** | **PAG network** | **PCG network** |
| --- | --- | --- |
| **Number of nodes** | 490 | 32 |
| **Number of edges** | 33061 | 449 |
| **Node degree exponent** | 0.128 | -4.125 |
| **Clustering coefficient** | 0.650 | 0.937 |
| **Connected components** | 15 | 1 |
| **Network diameter** | 6 | 2 |
| **Network centralization** | 0.335 | 0.101 |
| **Characteristic path length** | 1.886 | 1.095 |
| **Network heterogeneity** | 0.678 | 0.121 |
| **Network density** | 0.276 | 0.905 |

**Figure 3:**
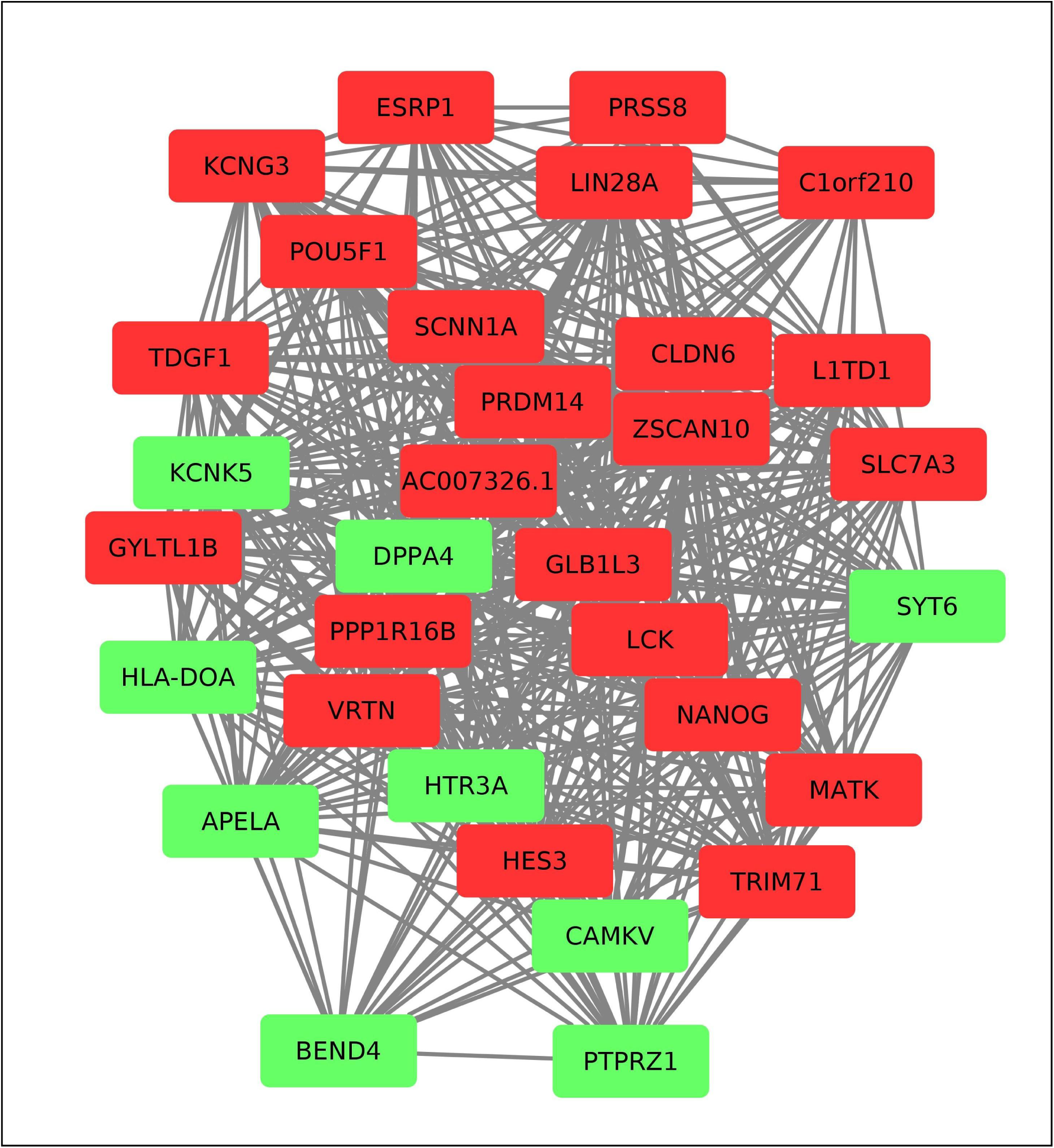
Pluripotency critical genes (PCGs) network. Co-expression network of the overexpressed genes in each datasets were reconstructed separately and then merged to obtain the PAG network. The PCG network consisting of the 32 PCGs was extracted from the PAG network. The different node colours here represent the cluster to which they grouped. Red for cluster-1 and green for cluster-2.

We used the community clustering algorithm to assess if the nodes of the major component have any natural tendency to group themselves. Two major clusters were formed consisting of 231 and 227 nodes. A third smaller cluster consisted of 10 nodes while the remaining eight nodes were found to be occurring as duplexes. Supplementary File 3 lists the genes in each cluster. The pluripotency critical genes were distributed among cluster-1 and cluster-2; with a majority (23 of 32 genes) occurring in cluster-1.

### 3.3. Enrichment analysis of co-expression network gene clusters

Enrichment analysis of the genes in the two major clusters were performed using PANTHER to identify if any classes of genes are over-represented by the gene sets. In cluster-1, 228 (out of 231) genes and in cluster-2, 222 (out of 227) genes mapped to the PANTHER database. The genes in cluster-1 were found to be enriched in biological processes terms related to developmental processes, system development, and cell-cell junction organization while the terms related to cell cycle, nuclear division and DNA replication were found to be enriched in cluster-2 (Supplementary File 4). The over-representation of these terms is a direct evidence towards the involvement of the genes in pluripotency. Further, when we checked if the genes of these significant terms make up any known biological pathway, we found 16 and 50 genes from cluster-1 and cluster-2 respectively mapped to the significant Reactome pathways (Supplementary File 4). Nearly 40% of the genes from each cluster were found to remain unclassified suggesting that more work has to be carried out in order to ascertain the pathways through which the genes function. Also, the wide gap in the number of genes that mapped to biological processes and pathways depict that though the individual function of a gene is known, the particular pathway through which it acts is unclear. The association of the genes together in the co-expression clusters in turn gives a positive indication of the biological processes and pathways the uncharacterised genes might be involved in paving a way for experimental biologist to plan their studies.

### 3.4. Validation of pluripotency related genes

In order to examine whether the genes identified through our study overlaps with previously established and curated stemness associated gene signatures we used StemChecker (Pinto et al., 2015). Among the 498 PAGs, 480 were found in the StemChecker database and 348 overlapped with atleast one stemness signature or transcription factor sets. On the other hand, 30 of the 32 PCGs were found in the StemChecker database and 25 genes overlapped with at least one stemness signature set (Supplementary File 5). In both the cases, there was significant overlap with ESC gene signatures. For the remaining unvalidated seven PCGs we used Expression Atlas (Petryszak et al., 2015) to individually review their expression levels in ESCs. We found at least one experiment for each gene that showed these genes to be upregulated in pluripotency. Further, on comparing our pluripotency related genes with the recently identified 50 gene hESC-essentialome, we found an overlap of 6 PCGs and 23 PAGs (Yilmaz et al., 2018).

### 3.5. Pluripotent cell differentiation markers

Parallel investigation of downregulated genes were carried out to identify genes that might serve as differentiation markers. We found 432 genes that were common across all the five datasets and detected by the three tools in each case (Table 2). Further, comparison of the top 10% downregulated genes revealed eight genes, namely DCN, FOXC1, GBP1, LUM, IGFBP7, COL3A1, HSPB3, and TMEM173, common across the datasets. Co-expression network of the downregulated genes were created in an identical manner as described for overexpressed genes in section 3.2. The merged network of the downregulated genes contained 386 nodes and 13,807 edges (Supplementary File 2). The network of the top 10% downregulated genes containing 8 nodes connected by 10 edges was extracted from the merged network and is shown in Figure 4. GO analysis of the 432 common downregulated genes showed the biological processes terms regulation of stem cell differentiation, extracellular matrix organization, anatommical structure morphogenesis, heart development, circulatory system development, tissue development, and regulation of fibroblast proliferation among the significantly enriched terms and clearly indicate the involvement of the genes in pluripotent cell differentiation. Similarly, most significantly enriched Reactome pathways were related to extracellular matrix (ECM) organization including ECM proteoglycans, elastic fibre formation, ECM interactions and collagen degradation. Terms related to signaling pathways like signaling by MET, signaling by PDGF and signaling by interleukins also occurred among the enriched Reactome pathways. Cells after exiting from pluripotent state organizes itself to form functional tissues and organs. The proper arrangement of cells is ensured by the ECM. Thus the occurrence of terms related to ECM organization confirms the role of these genes during differentiation of pluripotent cells to non-pluripotent cells. Apart from this, the enrichment of terms related to heart development and fibroblast proliferation were expected as the non-pluripotent samples in our study were either cardiomyocytes or fibroblast. The expression of the eight genes were checked on Expression Atlas (Petryszak et al., 2015) and Amazonia database (Assou et al., 2007) and were indeed found to be specific to non-pluripotent cells, thus validating them as potential differentiation markers.

**Figure 4:**
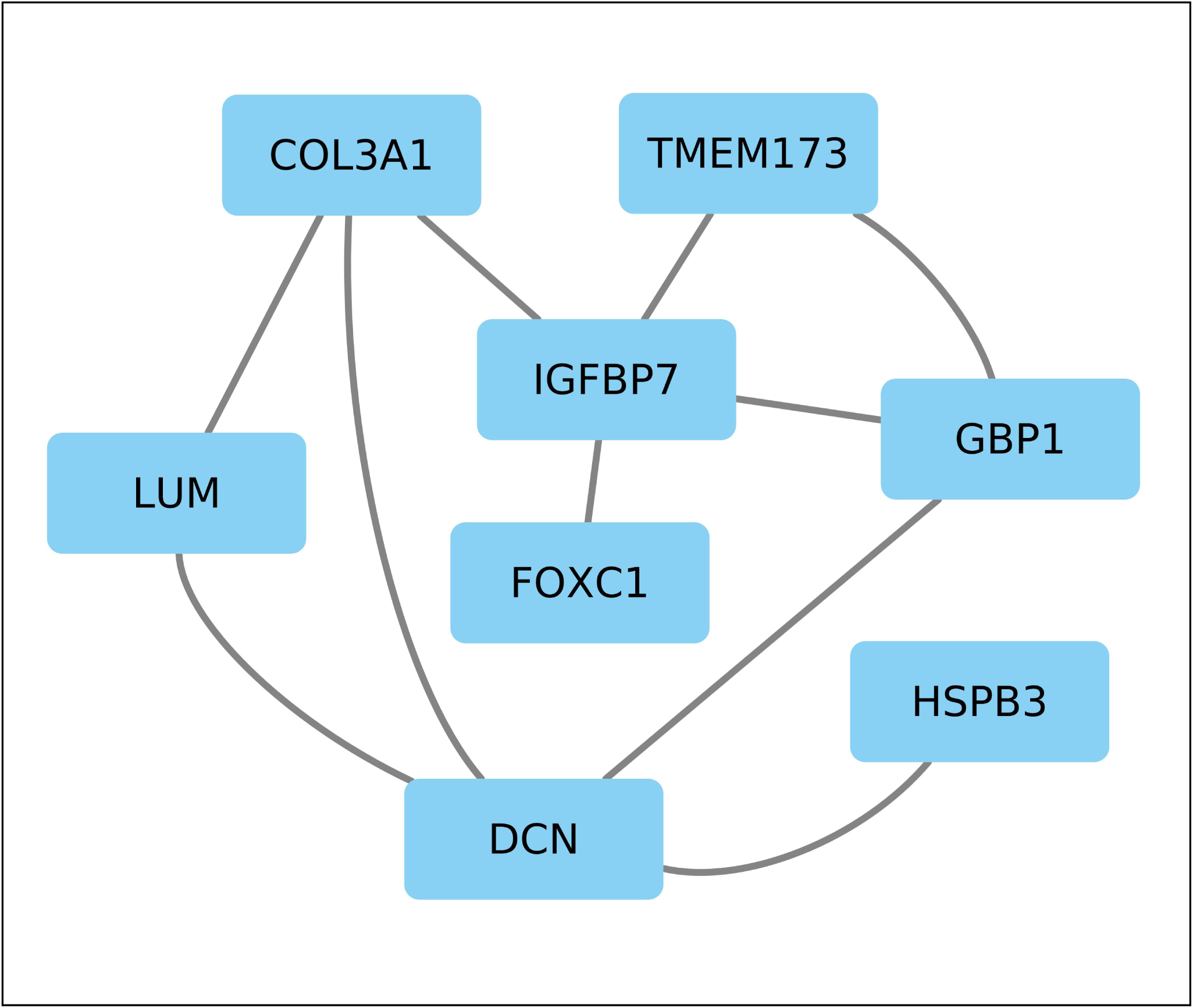
Network of the common top 10% downregulated genes.

## 4. DISCUSSIONS

We analysed transcriptomic data from five different RNA-seq experiments using a uniform pipeline. Our analysis revealed 32 genes (Table 4) to be critical for the establishment and maintenance of pluripotency. These 32 genes were among the top 10% overexpressed genes in all the five datasets and were also found to be co-expressed in all the datasets. Further, the network of the 32 genes exhibited a small world architecture depicting its importance. These 32 genes were divided into two groups based on the clustering of the co-expression network. The first 23 genes belong to cluster-1 that had an enrichment of genes involved in developmental processes, system development, and cell-cell junction organization. The remaining 9 genes belong to the cluster-2 and were enriched for terms related to cell cycle, nuclear division and DNA replication.

**Table 4:** List of pluripotency critical genes (PCGs)

| S. No<br>. | Ensembl Gene ID | Gene Name | Description |
| --- | --- | --- | --- |
| 1 | ENSG00000253313 | C1orf210 | Chromosome 1 open reading frame 210 |
| 2 | ENSG00000184697 | CLDN6 | Claudin 6 |
| 3 | ENSG00000104413 | ESRP1 | Epithelial splicing regulatory protein 1 |
| 4 | ENSG00000166105 | GLB1L3 | Galactosidase beta 1 like 3 |
| 5 | ENSG00000165905 | GYLTL1B | Glycosyltransferase-like 1B |
| 6 | ENSG00000173673 | HES3 | Hes family bHLH transcription factor 3 |
| 7 | ENSG00000171126 | KCNG3 | Potassium channel, voltage gated modifier subfamily G, member 3 |
| 8 | ENSG00000240563 | L1TD1 | LINE-1 type transposase domain containing 1 |
| 9 | ENSG00000182866 | LCK | LCK proto-oncogene, Src family tyrosine kinase |
| 10 | ENSG00000131914 | LIN28A | Lin-28 homolog A (C. elegans) |
| 11 | ENSG00000007264 | MATK | Megakaryocyte-associated tyrosine kinase |
| 12 | ENSG00000111704 | NANOG | Nanog homeobox |
| 13 | ENSG00000204531 | POU5F1 | POU class 5 homeobox 1 |
| 14 | ENSG00000101445 | PPP1R16B | Protein phosphatase 1 regulatory subunit 16B |
| 15 | ENSG00000147596 | PRDM14 | PR domain containing 14 |
| 16 | ENSG00000052344 | PRSS8 | Protease, serine 8 |
| 17 | ENSG00000111319 | SCNN1A | Sodium channel, non voltage gated 1 alpha subunit |
| 18 | ENSG00000165349 | SLC7A3 | Solute carrier family 7 (cationic amino acid transporter, y+ system), member 3 |
| 19 | ENSG00000241186 | TDGF1 | Teratocarcinoma-derived growth factor 1 |
| 20 | ENSG00000206557 | TRIM71 | Tripartite motif containing 71, E3 ubiquitin protein ligase |
| 21 | ENSG00000133980 | VRTN | Vertebrae development associated |
| 22 | ENSG00000130182 | ZSCAN10 | Zinc finger and SCAN domain containing 10 |
| 23 | ENSG00000279560 | AC007326.1 |  |
| 24 | ENSG00000248329 | APELA | Apelin receptor early endogenous ligand |
| 25 | ENSG00000188848 | BEND4 | BEN domain containing 4 |
| 26 | ENSG00000164076 | CAMKV | CaM kinase-like vesicle-associated |
| 27 | ENSG00000121570 | DPPA4 | Developmental pluripotency associated 4 |
| 28 | ENSG00000204252 | HLA-DOA | Major histocompatibility complex, class II, DO alpha |
| 29 | ENSG00000166736 | HTR3A | 5-hydroxytryptamine (serotonin) receptor 3A, ionotropic |
| 30 | ENSG00000164626 | KCNK5 | Potassium channel, two pore domain subfamily K, member 5 |
| 31 | ENSG00000106278 | PTPRZ1 | Protein tyrosine phosphatase, receptor-type, Z polypeptide 1 |
| 32 | ENSG00000134207 | SYT6 | Synaptotagmin 6 |

Our list of pluripotency critical genes includes some of the well-established pluripotency-associated transcription factors namely NANOG, POU5F1(OCT4), ZSCAN10, PRDM14 and HES3 (Hirata et al., 2000; Yu et al., 2009; Chia et al., 2010). NANOG prevents the differentiation of the hESCs to extraembryonic endoderm and trophectoderm lineages and hence acts as a gatekeeper of pluripotency (Hyslop et al., 2005). POU5F1 is a dose-dependent cell fate determinant and controls the expression of a number of genes involved in embryonic development (Niwa et al., 2000; Xu et al. 2009). ZSCAN10 is an ESC specific transcription factor that directly regulates POU5F1 expression by binding to its promoter (Yu et al., 2009). PRDM14 is another transcription factor that is known to influence the expression level of POU5F1. It directly upregulates POU5F1 expression through its proximal enhancer. PRDM14 has both positive and negative roles in transcription (Chia et al., 2010). HES3 is a transcriptional repressor of genes that require a bHLH (basic helix-loop-helix) protein for their transcription. Proteins transcribed by the Hes-family are known to regulate the binary cell fate decisions in many organs and the timing of several developmental events such as somite segmentation (Hirata et al., 2000).

Our list of critical genes for pluripotency included four RNA binding proteins (RBPs) namely LIN28A, TRIM71, L1TD1, and ESRP1. LIN28A and TRIM71 inhibit the processing of pre-let-7 miRNAs and regulates translation of mRNAs that control developmental timing, pluripotency, and metabolism (Rybak et al., 2009; Jin et al., 2011; Worringer et al., 2014). L1TD1 is a stem-cell specific RBP and is responsible for the post-translational nuclear import of the core pluripotency transcription factors OCT4, SOX2 and NANOG from cytoplasm. ESRP1 has been found to help fine-tune the expression of pluripotency related factors and maintain balance between its property of self-renewal and commitment to restricted developmental fate (Fagoonee et al., 2013).

Four genes, namely SCNN1A, KCNG3, KCNK5, and HTR3A, involved in the formation of ion channels can be found in our pluripotency critical gene list. Ion channels play a crucial role in signal transduction and its importance is well known in embryonic development (Tosti et al, 2016). Previous studies have reported that proliferation of hESCs is inhibited when ion channels are blocked (Wang et al., 2005). SCNN1A is a sodium permeable non-voltage sensitive ion channel protein while KCNG3 and KCNK5 are potassium channel proteins. HTR3A is a ligand-gated ion channel protein and one of the several receptors for serotonin; a biogenic hormone that functions as a mitogen (Fanburg and Lee, 1997).

LCK is a proto-oncogene belonging to the Src-family of non-receptor tyrosine kinases (nRTKs). Members of these family of kinases are known to be coupled to many growth factor receptors to regulate cell adhesion, proliferation, growth and survival (Zhang et al., 2014). In a study that aimed to monitor the expression of this protein during embryoid body formation found that the transcript and protein levels were lost more rapidly than that of the core pluripotency transcription factor POU5F1. The study further made an observation that the changes in LCK expression and activity are unique to hESCs against mouse ESCs suggesting a possible role in maintenance of undifferentiated state. Another nRTK, MATK, belonging to the Csk-family can be found in our gene list. The Csk-family of nRTKs are known to inhibit the tyrosine kinase activity of Src-family kinases (Okada, 2012).

DPPA4 is an oncogene that functions independently of the core pluripotency network (Klein et al., 2018). APELA functions as a regulatory RNA and promotes self-renewal via progression of cell cycle and protein translation. Inhibition of APELA has been shown to cause reduced hESC growth, cell death and loss of pluripotency (Ho et al., 2015).

Apart from these, we found several other genes known to be important for pluripotency but did not occur in our list of critical genes. However, they were upregulated and co-expressed in all the five datasets and also formed the major connected component in the merged network. These includes some of the crucial transcription factors namely ZIC3, FOXD3, SALL4, and SOX group proteins SOX3 and SOX13. ZIC3 functions as a direct activator of NANOG (Lim et al., 2010) while SALL4 regulates the transcription of POU5F1 (Zhang et al., 2006). Stem cells lacking FOXD3 maintained normal proliferation rate but showed an increased tendency to apoptosis (Liu and Labosky, 2008). SOX3 is a close relative of SOX2 and it has been accounted to be functionally equivalent to SOX2 (Adikusuma et al., 2017).

## 5. CONCLUSIONS

Comparison of transcriptomic data from pluripotent and non-pluripotent samples revealed 498 pluripotency associated genes (PAGs) and 432 genes as potential pluripotent cell differentiation markers. Further refinement of the PAGs led to 32 genes to be critical for the induction and maintenance of pluripotency in human. These pluripotency critical genes formed a tightly bound co-expression network with small-world architecture. Assessment of the pluripotency critical genes brought out the presence of some of the well-known pluripotency related genes like NANOG, POU5F1, ZSCAN10, LIN28, L1TD1, etc. Apart from these, there were several other genes like SCNN1A, SLC7A3, MATK, PRSS8, etc. that have been linked to pluripotency but their exact molecular function and underlying mechanisms are yet to be elucidated in human. Thus more experimental as well as theoretical works are necessary towards unravelling the pluripotency players and their interaction/regulation network which is indispensable for the successful applications of hESCs.

## ACKNOWLEDGEMENTS

AS thank the Department of Biotechnology (DBT) for providing financial assistance for the project. Thanks to Ms. Rinki Singh, Ms. Priyanka Kumari and Mr. Amresh Sharma for their helpful discussions.

## SUPPLEMENTARY DATA

**Supplementary File 1:** Details of raw data included in the study.

**Supplementary File 2:** Cytoscape file of the gene co-expression networks. The network names are as follows:

PAG Network : Pluripotency associated gene network.

PAG Network - CC1 : The component of the PAG network consisting of majority of the nodes.

PCG Network : Network of pluripotency critical genes.

PAG Network - CC1--clustered GLay : The result of clustering the larger component of the PAG network.

PAG Network - CC1--clustered GLay(C1/2/3) : Sub-networks of Clusters 1/2/3.

Downreg merged network: The network formed by merging the individual co-expression networks of the downregulated genes in each datasets.

Top10pc downreg network: Network of the common top 10% downregulated genes.

**Supplementary File 3:** List of genes in each cluster of the PAG network.

**Supplementary File 4:** Results of the functional and pathway analysis of genes of two major clusters in the PAG network.

**Supplementary File 5:** StemChecker analysis results of pluripotency critical genes (PCG).

