## Supplementary File 5 for "RNA-Seq analysis reveals pluripotency-associated genes and their interaction networks in human embryonic stem cells"

**Supplementary File 5:** StemChecker analysis results of pluripotency critical genes (PCG).

StemChecker was used to validate the 32 genes that were found to be critical for pluripotency from our analysis. Of the 32 genes, 25 genes could be mapped to at least one stemness signature. The checkerboard table below displays the occurrence of queried genes across the stemness signatures. Blue squares indicate the presence of the gene in the corresponding stemness signature set.

| Stemness signature | Hs_ESC_Assou | Hs_ESC_NANOG_targets_Boyer | Hs_ESC/EC_Sperger | Hs_iPSC_Shats | Hs_ESC_SOX2_targets_Boyer | Hs_ESC_Skottman | REACTOME | Plurinet | Hs_ESC_OCT4_targets_Boyer | Hs_EC_Skotheim | GeneCards | Hs_ESC_Chia | Hs_ESC_Bhattacharya | Hs_ESC_Sato | Hs_HSC_Huang | Hs_HSC_Novershtern | KEGG | Hs_NSC_Huang | Hs_HSC_Toren |
|---|---|---|---|---|---|---|---|---|---|---|---|---|---|---|---|---|---|---|---|
| Gene Names | | | | | | | | | | | | | | | | | | | |
| POU5F1 | ■ | ■ | ■ | ■ | ■ | ■ | ■ | ■ | ■ | ■ | ■ | ■ | ■ | ■ | | | ■ | | |
| NANOG | ■ | ■ | | ■ | ■ | ■ | ■ | ■ | ■ | ■ | | | ■ | | | ■ | ■ | | |
| DPPA4 | ■ | ■ | ■ | | 1 | 1 | | | ■ | ■ | | | | ■ | ■ | ■ | | | ■ |
| TDGF1 | ■ | ■ | | ■ | ■ | ■ | ■ | ■ | ■ | | | | ■ | | | | | | |
| PRDM14 | ■ | ■ | | | ■ | | ■ | | ■ | ■ | ■ | ■ | | | | | | | |
| LIN28 | ■ | | | ■ | | ■ | ■ | | | | ■ | | | | | | | | |
| SCNN1A | ■ | ■ | ■ | ■ | ■ | | | | | | | | | | | | | | |
| FLJ20171 (ESRP1) | ■ | | ■ | ■ | | ■ | | | | | | | | | | | | | |
| ZSCAN10 | ■ | ■ | | | | | ■ | | | | | ■ | | | | | | | |
| LCK | ■ | | | ■ | | | | ■ | | | | | | | | | | | |
| PTPRZ1 | ■ | | ■ | | | | | | | | | | | | | | | ■ | |
| C14orf115 (VRTN) | ■ | ■ | | | ■ | | | | | | | | | | | | | | |
| PPP1R16B | ■ | ■ | | ■ | | | | | | | | | | | | | | | |
| PRSS8 | | ■ | ■ | | ■ | | | | | | | | | | | | | | |
| TRIM71 | | | | | | ■ | | | | | ■ | | | | | | | | |
| SYT6 | ■ | | ■ | | | | | | | | | | | | | | | | |
| MATK | ■ | | | | | | | ■ | | | | | | | | | | | |
| FLJ10884 (L1TD1) | ■ | | ■ | | | | | | | | | | | | | | | | |
| CLDN6 | ■ | ■ | | | | | | | | | | | | | | | | | |
| BEND4 | | | | | | | | | | | | | | | ■ | | | | |
| KCNK5 | ■ | | | | | | | | | | | | | | | | | | |
| Hs.105196 (APELA) | ■ | | | | | | | | | | | | | | | | | | |
| SLC7A3 | ■ | | | | | | | | | | | | | | | | | | |
| MGC8407 (CAMKV) | ■ | | | | | | | | | | | | | | | | | | |
| GYLTL1B | ■ | | | | | | | | | | | | | | | | | | |

| Colour code for source type | |
|---|---|
| | TF Target Genes |
| | Expression Profiles |
| | RNAi Screens |
| | Literature Curation |
| | Computationally Derived |

### Source datasets:

7. **REACTOME:** Croft, D., Mundo, A.F., Haw, R., Milacic, M., Weiser, J., Wu, G., Caudy, M., Garapati, P., Gillespie, M., Kamdar, M.R. and Jassal, B., 2013. The Reactome pathway knowledgebase. Nucleic acids research, 42(D1), pp.D472-D477.

8. **Plurinet:** Müller, F.J., Laurent, L.C., Kostka, D., Ulitsky, I., Williams, R., Lu, C., Park, I.H., Rao, M.S., Shamir, R., Schwartz, P.H. and Schmidt, N.O., 2008. Regulatory networks define phenotypic classes of human stem cell lines. Nature, 455(7211), p.401.

9. **Hs_ESC_OCT4_targets_Boyer:** Boyer, L.A., Lee, T.I., Cole, M.F., Johnstone, S.E., Levine, S.S., Zucker, J.P., Guenther, M.G., Kumar, R.M., Murray, H.L., Jenner, R.G. and Gifford, D.K., 2005. Core transcriptional regulatory circuitry in human embryonic stem cells. cell, 122(6), pp.947-956.

10. **Hs_EC_Skotheim:** Skotheim, R.I., Lind, G.E., Monni, O., Nesland, J.M., Abeler, V.M., Fosså, S.D., Duale, N., Brunborg, G., Kallioniemi, O., Andrews, P.W. and Lothe, R.A., 2005. Differentiation of human embryonal carcinomas in vitro and in vivo reveals expression profiles relevant to normal development. Cancer research, 65(13), pp.5588-5598.

11. **GeneCards:** Safran, M., Dalah, I., Alexander, J., Rosen, N., Iny Stein, T., Shmoish, M., Nativ, N., Bahir, I., Doniger, T., Krug, H. and Sirota-Madi, A., 2010. GeneCards Version 3: the human gene integrator. Database, 2010.
